# A Paradigm for Measuring Resting State Functional Connectivity in Young Children Using fNIRS and Freeplay

**DOI:** 10.1101/2020.01.13.904029

**Authors:** Jaeah Kim, Alexander Ruesch, Nin Rebecca Kang, Theodore J. Huppert, Jana Kainerstorfer, Erik D. Thiessen, Anna V. Fisher

**Author notes:** **Corresponding author:** Jaeah Kim.

## Abstract

Resting state functional connectivity (RSFC) reflects the organization of functional networks in the brain. Functional networks measured during “resting”, or task-absent, state are correlated with cognitive function, and much development of these networks occurs between infancy and adulthood. However, RSFC research in the intermediate years (especially between ages 3 and 5 years) has been limited, mainly due to a paucity of child-appropriate neural measures and behavioral paradigms. This paper presents a new paradigm to measure RSFC in young children, utilizing functional near-infrared spectroscopy (fNIRS) and Freeplay, a simple behavioral setup designed to approximate resting state in children. In Experiment 1, we recorded fNIRS data from children aged 3-8 years and adults aged 18-21 years and examined feasibility and validity of our measure of RSFC, and compared measures across the two groups. In Experiment 2, we recorded longitudinal data at two points (approximately 3 months apart) from children aged 3-5 years, and examined reliability under a variety of measures. In both experiments, all children were able to complete testing and provide usable data, a significant improvement over fMRI-based RSFC measurement in children. Results suggest this paradigm is practical and has good construct validity and test-retest reliability, and may contribute towards increasing the availability of reliable data on resting state networks in early childhood. In particular, these are some of the first positive results on the feasibility of reliably measuring functional connectivity in children aged 3-5 years.

## Introduction

Brain function arises from a concerted effort of various regions working together in what can be characterized as networks. Research in the past few decades has focused on properties of these networks and patterns of connectivity that support cognitive functions [1–3]. Resting state networks (RSNs) -- networks of regions showing temporal correlation of low-frequency fluctuations (functional connections) in subjects not performing any task [4] -- can reflect underlying functional organization of the brain.

Discovery of resting state functional connectivity (RSFC) and its link to function [5–8] has widened the door to investigating mechanisms of cognitive function by providing some advantages over task-based studies: 1) whereas measuring functional connectivity during task provides us with the connectivity of only those regions specifically involved in the task, measurement during resting state can provide connectivity information simultaneously among many regions; 2) certain populations (e.g., clinical populations) or cognitive functions (e.g., motor) that are difficult to engage with specific tasks in the scanner, can be just as well studied in resting state. For these reasons, resting state measurement has become a standard paradigm for studying functional connectivity in human adults.

### Measuring RSN development

RSN research in infants and adults presents a clear developmental trend: whereas 10-20 RSNs have been identified in adults, only about 5 of these networks are found in infants, the rest likely developing throughout childhood [9,10]. This suggests that the functional architecture of the brain may be undergoing dramatic development and restructuring in these intermediate years to create many of these networks that emerge prominently by adulthood. *Patterns* of connectivity also change over development-- a number of studies have shown that children display more diffuse functional connectivity patterns and increased connectivity with adjacent regions, while adults show more focal connectivity patterns and increased connectivity between distant regions [11,12]. This simultaneous increase in both segregation (pruning short-range connections) and integration (strengthening long-range connections) of brain regions over development likely reflects a transition from organization around spatial proximity to organization around higher-order function [12]. Finally, aberrant connectivity in RSNs have been associated with a variety of psychopathologies from affective disorders such as depression, to diseases of cognitive function such as Alzheimer’s disease, to neurodevelopmental disorders such as autism or ADHD (for review, see [10]), suggesting that RSN development may be integral to healthy brain and cognitive development. Together, these findings highlight the need for studying RSNs over the time course of development and especially in early childhood, when perhaps the most change is occurring in some brain networks.

However, studying RSNs in children has been challenging, mainly for two reasons. First, traditional neuroimaging tools (e.g., functional MRI (fMRI), electroencephalography (EEG), or magnetoencephalography (MEG)) are difficult to utilize with awake children [13,14]. Second, the standard procedure for measuring resting state connectivity -- to sit still for a period of time -- proves to be a difficult task for children, and some methods previously employed to increase compliance in children during resting state measurement, such as movie or video-watching, have been shown to significantly alter resting state in adult participants [15]. Consequently, resting state studies in children and especially in young (3-5 years/pre-school aged) children have been limited [16]. The current study uses functional near-infrared spectroscopy (fNIRS) which provides an appropriate solution to child neuroimaging challenges and introduces a new paradigm called Freeplay to address the difficulty of employing an appropriate “task” for resting state measurement – the combination provides a simple and efficient paradigm to improve participant compliance and quality RSFC measurement (of surface regions) in children. By reconstructing characteristic features of RSFC, we demonstrate the feasibility of this fNIRS-Freeplay set-up for studying resting state connectivity in preschool and early school-aged children. We also study the test-retest reliability of the paradigm under a number of measures, showing that consistent results can be derived across multiple scans, at both individual and group levels. Some multi-session test-retest reliability studies for RSFC measurement methods exist for adults and older children in fMRI, as well as for adults in fNIRS, but not for children in fNIRS [17–20].

### fNIRS with young children

Traditionally studied with fMRI, and to a lesser degree with EEG/MEG, RSNs have been studied primarily in adult and sleeping infant populations, but relatively rarely in child populations, as these imaging techniques are challenging to use with awake children [21]. For example, a major difficulty is the sensitivity of these modalities to movement artifacts combined with children’s difficulty remaining still for extended periods of time, which can systematically bias functional connectivity measures [22]. In a previous study on the feasibility of fMRI measurement in children, Byars et al. [13] reported a 47% success rate with children between the ages of 5 and 6, under a relatively generous definition of “success” as ‘completing at least one of four fMRI tasks and an anatomical reference scan’ [14] (p. 2). Moreover, another more recent study of fMRI feasibility with children and adolescents showed that clinical groups scanned even less successfully than typically developing controls [14].

fNIRS is a relatively recent light-based neuroimaging method that overcomes many of the challenges with obtaining brain activity measures in child populations. In fNIRS, near-infrared light is used to obtain an estimate of changes in both oxygenated and deoxygenated hemoglobin concentrations in a region of the brain. Thus, like fMRI, fNIRS gives an indirect measure of neural activity based on blood oxygenation levels. Compared to fMRI or EEG, fNIRS is robust to and unrestrictive of motion, comfortable, quick to set up, and cost effective (see [23] for comparison of techniques). These factors make it especially appropriate for use with children. The main limitation of fNIRS in the context of studying RSFC is that measurement is limited to regions near the surface of the brain (<17 mm of brain tissue deep). For researchers interested in studying RSFC among surface regions of the brain, this is a viable neuro-measurement tool. Our study, as discussed below, measures from the surface of the prefrontal cortex.

### Freeplay in a fNIRS recording set-up can overcome challenges in behavioral methods

Resting state studies have mainly recorded from adults instructed to remain still for a period of time [24]. Some RSFC studies are conducted in infants during sleep [25]. Unfortunately, neither recording situation is practical for young children; children often have great difficulty remaining quietly still for an extended time, and unlike infants, children are far less susceptible to maintaining sleep in the scanner. Moreover, it is not clear that sleep is a good approximation of resting state, as it exhibits its own distinctive functional connectivity patterns [26–29]. For such reasons, resting state-fMRI studies that are run with children frequently involve extensive and costly effort in pre-training children in “mock” scanner environments to help reduce motion artifacts and improve participant compliance. In spite of this pre-training, unwieldy proportions of scans (e.g., ranging from as little as 18% to as much as 53% in children aged 4-6) are discarded due to movement artifacts contaminating the signal [30,31].

To mitigate this, many resting state-fMRI studies with children and patient populations have used motion picture stimuli, with or without audio, to help engage participants in the fMRI scanner. These have had some success in improving participant compliance (e.g., reduced participant motion and reduced frequency of falling asleep) [32,33]. However, viewing movie clips have been found to significantly alter resting state in adult participants [15]. Furthermore, viewing movie clips may engage task-specific networks, such as those involved in language or audition. In fact, movies are sometimes used in studies as stimuli or tasks [34], or have specifically been reported to induce different connectivity patterns compared to rest (e.g., early visual network decreased its connectivity with dorsal attention network and increased its connectivity with the default mode network as well as the fronto-parietal network, during movie watching [33]. These findings bring into question the validity of using movies for studying “resting state” -functional connectivities and -networks in particular. However, movie or video-watching still remains the most feasible method by which to record RSFC from children in an fMRI setting (with varying success depending on where the movie lies in the spectrum from too engaging to not engaging enough).

Because fNIRS recording set-up is quiet, comfortable to wear, and unrestrictive compared to the fMRI environment, a less engaging set-up for aiding compliance can be sufficient. Our study proposes an experimental paradigm called Freeplay that may closely approximate resting state, and which takes advantage of the spatially unrestrictive nature of an fNIRS recording situation. In this “task”, participants are seated at a table, presented with a set of simple toys (e.g., wooden blocks, small plastic animals) and asked to quietly play for a few minutes. The premise is that children can naturally comply much more easily with sitting still and quietly for a period of time when presented with even simple and unengaging toys. In addition, the fNIRS possible sampling rate is much greater than that of fMRI, thus requiring less recording time overall. Due to the simple nature of the toys, Freeplay is expected to induce quiet boredom, a state we expect may closely approximate resting state. Additionally, the unconstrained nature of this set-up naturally mirrors the “at rest” eyes-open set-up for adults in which adults sit open-endedly for a period of time without any specific task, and in the same way, allows natural individual variation to be present. Allowing natural variability is important as it allows researchers to be more confident that connectivity patterns consistent across many participants are generalizable and not arising from task-specific or stimuli-specific states.

Since measuring RSFC in children relies on using *approximations* of the resting state task (e.g., movie watching, silent rest), it is important to also develop multiple such approximations, to combat the task impurity problem. Freeplay contributes as one method for approximating resting state. Moreover, Freeplay avoids several biases that are inherent in some previous methods. For example, relatively engaging or externally-guided stimuli, such as movie clips or screensavers, potentially compromise the unrestrained and non-externally-directed nature of thinking that is characteristic of true resting state. They may also potentially engage functional networks differently from rest (e.g., naturalistic movie viewing has been shown to alter connectivity patterns among certain networks compared to rest) [33]. In contrast, Freeplay’s undirected or internally-/self-guided nature may more closely mirror the state of adults in resting state. Additionally, in contrast to other proposed methods that utilize identical or similar visual stimuli sequences across participants (movies, screensaver-type stimuli, e.g., Inscapes), Freeplay lacks any time-locking events that could introduce systematic biases when aggregating data across subjects and artificially inflate across-subject consistencies [35].

### Prefrontal Cortex

The prefrontal cortex (PFC) houses major components of such RSNs as the central executive network (CEN) and default mode network (DMN). Its development, from infancy through adulthood, is prominently linked to the development of executive function (EF) [36], a collective system of basic cognitive processes that includes inhibition, working memory, and cognitive flexibility, and supports higher-order processes such as planning and problem solving [37,38]. PFC’s protracted development make it an especially interesting region to study over age, including in early development. Aberrant functional connections within the PFC have also been associated with ADHD symptoms and impaired inhibitory and attentional control, implicating its important role in health executive function development [39]. Additional advantages of measuring PFC with fNIRS are its close proximity to the surface of the skull and convenient placement under the forehead (which lacks hair, improving fNIRS signal quality). Thus, we focused data collection on the PFC in this validation study.

### Current study

This study aims to demonstrate feasibility of using fNIRS and Freeplay to measure RSFC in pre-school-aged and early school-aged children. We present this paradigm as a potential means to address the gap in research on RSFC in children, stemming from a lack of appropriate measurement and behavioral tools for the population. This set-up is designed to place minimal restrictions on the participant, allow sufficient data collection, and achieve relatively good signal-to-noise ratio (by minimizing sources of noise introduced by the measurement tool as well as the participant). This study aims to establish fundamental psychometric properties of the fNIRS-Freeplay paradigm, including construct validity, test-retest reliability, and feasibility.

## Experiment 1: Comparison with traditional adult resting state

We first investigated whether the fNIRS-Freeplay paradigm allows us to measure RSFC -- specifically, whether the paradigm exhibits construct validity. We did this in two ways. First, we asked whether the fNIRS-Freeplay paradigm reproduces a characteristic feature of adult resting state connectivity, namely strong connectivity between homologous (bilaterally symmetric) regions of the two hemispheres [40,41]. Second, we studied the distinguishability of RSFC in adults in Freeplay and adults in true resting state, in terms of the ability of machine learning classifiers to correctly classify instances of each.

If adults “at rest” and in Freeplay show *similar* connectivity patterns and are difficult to distinguish from each other, this would suggest that Freeplay may be a good approximation of resting state. Confirming this hypothesis would help validate fNIRS-Freeplay as a paradigm for measuring RSFC in adults, and a natural next step would be to apply this paradigm to measure RSFC in children. To begin exploring this (and also to provide a control for our first similarity measure -- classification error between adults in Freeplay and “at rest”), we additionally measured NIRS-Freeplay data in children, hypothesizing that adults and children in Freeplay will show *different* connectivity patterns, consistent with research suggesting significant development of RSFC from childhood into adulthood [9].

### Methods

#### Participants

Participants were 13 undergraduates (aged 18-21) from Carnegie Mellon University (CMU) and 18 children (aged 3 to 8 years, *M*_age_ = 4.8, *Med*_age_ = 4.3) recruited from the community and the Children’s School, a CMU-affiliated laboratory school. 17 children were included in the analysis after 1 exclusion due to experimenter error. Adults participated in both the standard resting state task and the Freeplay task within a single session. Task order was randomized. Children participated in Freeplay only.

#### Standard Resting State Task

Participants sat still and quietly at a desk with eyes open for 8 minutes. The scanning duration of 8 minutes sits comfortably over a 7-minute minimum reported to achieve accurate and stable RSFC from fNIRS measurement in children (aged 6.9 to 8.21 years) in a recent study by Wang, Dong, and Niu testing fNIRS RSFC test-retest reliability [42].

#### Freeplay Set-up

Participants sat quietly and freely played with a set of toys for about 8 minutes. Toys included: lincoln logs, wooden nuts and bolts, plastic animal figurines, toy cars, and simple coloring pages (a flower, turtle, duck, or fish). Toys were chosen to be simple and minimally engaging, to help induce quiet boredom, a state that we expect may closely approximate resting state (See Figure 1, <u>right panel</u>).

**Fig. 1.**
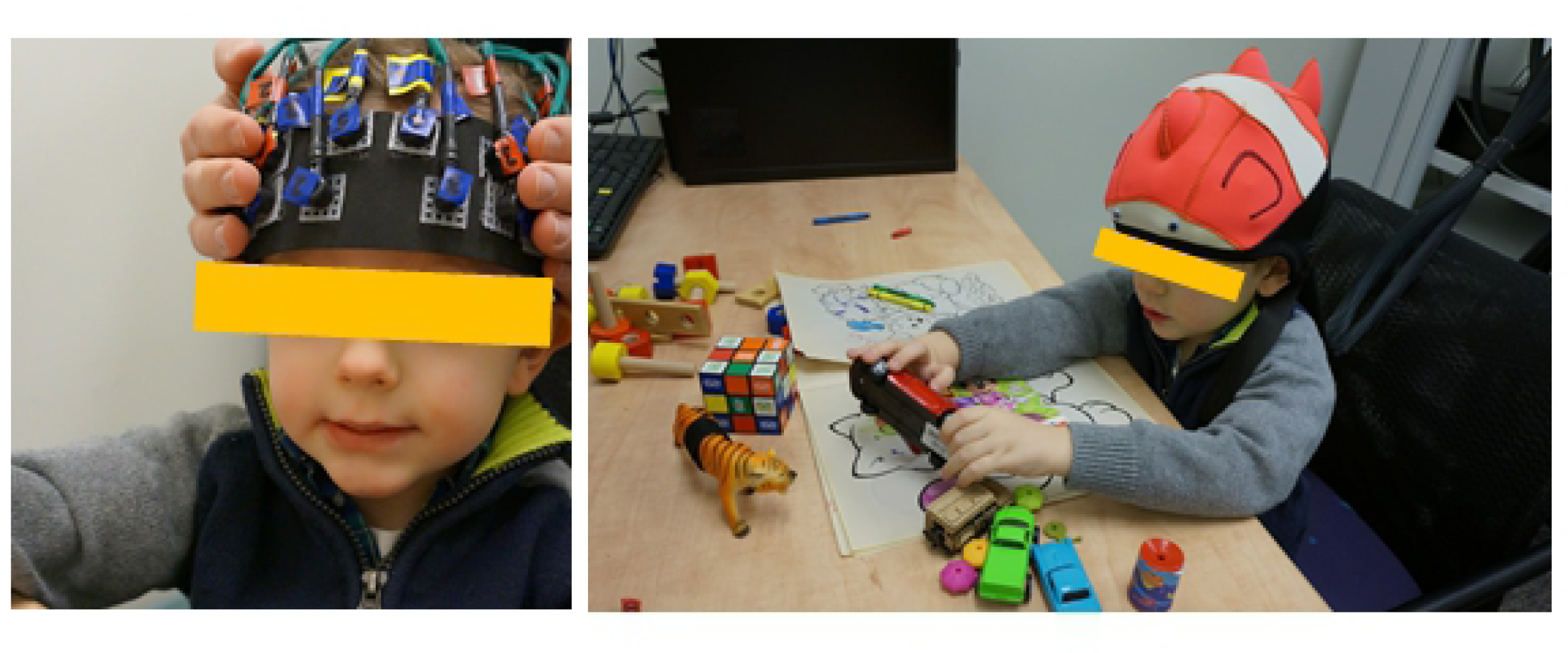
Placement of NIRS probe. The left picture displays NIRS probe strip placed over the forehead; the right picture shows probe strip secured by scuba cap and the freeplay setup with simple toys.

#### fNIRS Set-up

Neural activity was recorded at 20 Hz using a continuous wave real-time fNIRS system (CW6, Techen, Inc., Milford, MA, USA) with 4 light sources, each containing 690-nm (12 mW) and 830-nm (8 mW) laser light, and 8 detectors, to give oxy-hemoglobin and deoxy-hemoglobin measures in 10 channels on the PFC (Figure 1, left panel). Sensors were arranged in a layout as depicted in Figure 2. The distance between light sources and detectors were between 2.8 and 3 cm. Sensors were snapped into a cap strip built from foam sheet and plastic mesh, and connected to the fNIRS system via optic fibers. For each participant, the cap strip was positioned on the head, centered on position FpZ according to the international 10-20 coordinate system standard, extending over the Brodmann area 10 (anterior PFC) and area 46 (dorsolateral PFC) bilaterally. The strip was secured to the head using a neoprene scuba cap (pictured in Figure 1), to prevent probe from slipping as well as to cover the probe to prevent ambient light from reaching the sensors. The participant sat in a rigid, stationary chair to reduce movement artifacts.

**Fig. 2.**
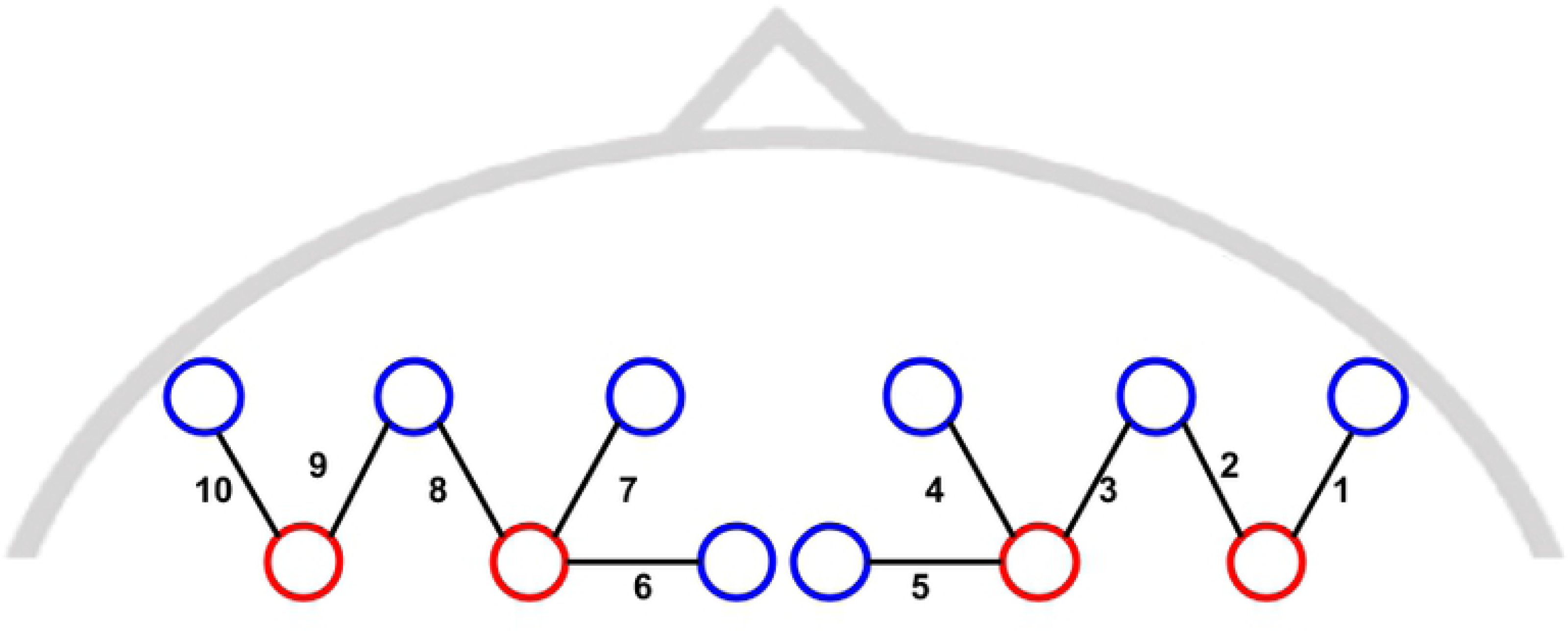
Probe layout for Experiment 1. Sources are in red and detectors are in blue. Channels are in black, labeled 1-5 on the right hemisphere and 6-10 on the left.

After fitting the fNIRS cap to the participant’s head, signal quality was checked for each source-detector channel and calibrated if needed to make sure the fNIRS fiber optics made good contact with the scalp of the participant, and that the detector was sensitive to cardiac pulsation as a sign of good signal-to-noise (SNR) ratio. Any detector saturation was also adjusted for in this step. fNIRS data was recorded for each participant using custom data collection software that interfaced with the fNIRS system, described in [43].

#### fNIRS Data Processing

Raw light attenuation measurements were converted to oxy-hemoglobin and deoxy-hemoglobin concentration changes using the modified Beer-Lambert law [44,45]. We removed long-term drifts in the data by subtracting a least-squares linear fit. We then band-pass filtered the data to remove cardiac and respiration signals (retaining frequencies in the range 0.01-0.1 Hz, as suggested by [40]). To mitigate motion artifacts in the form of sudden spikes or shifts, we applied a widely-used correlation based signal improvement (CBSI) filter, which is based on the assumption that true oxy-Hb and deoxy-Hb should be maximally negatively correlated [46]. As an alternative, we also ran all of the analyses on data filtered with a kurtosis-based wavelet filter (kbWF), shown previously to outperform other motion artifact removal methods such as PCA, tPCA, regular wavelet filter, and spline interpolation [47] -- results with the kbWF were qualitatively identical and hence not reported here. With the resulting time series, we computed partial correlations for each channel pair (CP), given the other channels (since there were 10 channels, there were 45 (10 choose 2) distinct CPs). These 45 computed partial correlations, which are represented graphically in correlation matrices (as in Figure 3) were the main quantities studied in this paper. We used partial correlation as the index of RSFC in this study because it can factor out correlation between fNIRS channels due to shared extracerebral components, and is thus thought to characterize relationships between brain regions more precisely than Pearson’s correlation [48,49].

**Fig. 3.**
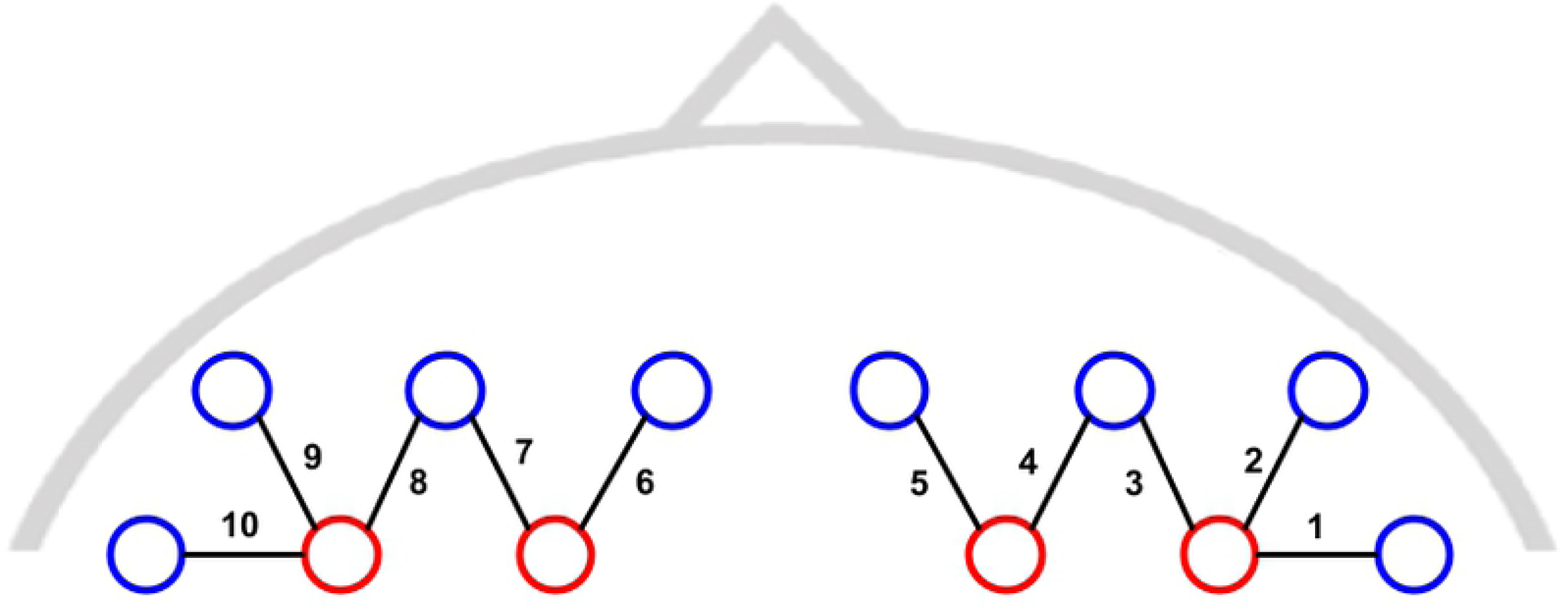
Group-averaged partial correlation matrices for children in Freeplay and for adults in Freeplay and at rest. Channels 1-5 were located on the right hemisphere; channels 6-10 were located on the left hemisphere. Homologous CPs are circled in the child panel.

#### Data Analysis Strategy

Our first goal was to test for significant homologous connectivity, characteristic of RSFC. To do this, we compared functional connectivity between regions that were homologous (bilaterally symmetric) to that between non-homologous regions.

Our second goal was to test the validity of Freeplay as a task for measuring resting state. To do so, we compared the functional connectivity in adults between the two conditions, “at rest” and Freeplay. Since it is difficult to directly measure similarity between two groups of functional connectivity patterns, we did so by estimating the accuracy of classifiers trained to distinguish different conditions (e.g., between “at rest” and Freeplay in adults). Higher accuracies suggest greater distinguishability, and hence greater dissimilarity, between classes. Given the high-dimensionality (45 CPs) of our problem, we used logistic LASSO (i.e., logistic regression with the Least Absolute Selection and Shrinkage Operator penalty), which should perform relatively well in our high-dimensional setting [50]. Accuracy was estimated using leave-one-out cross-validation (LOOCV). Within each LOOCV fold, the LASSO regularization parameter λ was selected by 10-fold cross validation. To reduce the chance that results were classifier specific, we also tried a highly distinct classifier, k-nearest neighbors (kNN) classification. Accuracy was again estimated by LOOCV, with k selected within each LOOCV fold by 10-fold cross-validation (over k=1,…,10, covering a range of common values [51]). To provide a baseline for comparison, we similarly compared data from adults versus children in Freeplay.

### Results

First and foremost, all participants (including all child participants) completed the task and provided usable data.

#### Homologous versus non-homologous CPs

First, we compared all homologous to all non-homologous pairs of distinct channels, to identify the strong inter-hemispheric homologous connections. Specifically, we averaged homologous and non-homologous CPs within subjects, forming two sets of measured CP correlations (*r*-values; separately for adults and children). This comparison showed that homologous CPs were significantly more strongly connected than non-homologous CPs (*p*<0.001 for both children and adults, by a two-sample *t*-test of the Fisher *z*-transformed *r*-values as well as a permutation test).

#### Adult correlations in Freeplay versus “at rest”

Next, we compared adult correlations in Freeplay and “at rest” to see how Freeplay compares with traditional resting state. Logistic LASSO, trained to predict Freeplay or “at rest” from CP correlations, gave a LOOCV accuracy of 38.462% (worse than chance, 50%), suggesting that the two tasks do not seem to elicit highly distinct connectivity (95% Wilson score confidence interval (CI): [19.76%, 57.16%]). The kNN classification to predict Freeplay or “at rest” from CP correlations achieved a LOOCV classification accuracy of 57.7% (95% Wilson score confidence interval (CI): [48.32%, 67.08%]), just over chance.

#### Adult versus child correlations in Freeplay

Next, we compared correlation matrices between adults and children, both in Freeplay. Logistic LASSO, trained to predict “adult” or “child” from CP correlations, gave a LOOCV accuracy of 83.333%, suggesting that adults and children exhibit highly distinct connectivity in Freeplay (95% Wilson score confidence interval (CI): [69.997%, 96.669%]).

## Experiment 2: Test-retest reliability with children in Freeplay

In this experiment, we studied reliability of the NIRS-Freeplay RSFC measure in children, in terms of consistency of results across independent scans (intersession, or test-retest, reliability). To do this, we collected multiple NIRS-Freeplay scans from children on different days and estimate a connectivity network from each scan. We then measured similarity of this estimated connectivity network both across scans within subjects and across subjects.

### Methods

#### Participants

Participants consisted of 19 children (aged 3 to 5 years, *M*_age_ during first scan = 4.34, *M*_age_ during second scan = 4.65, *Med*_age_ during first scan = 4.31, *M*_age_ during second scan = 4.82) recruited from the Children’s School. Data was collected longitudinally at 2 time points (on average 3.7 months apart), with 2 scans (approximately 1 week apart) at each time point. These data were collected as part of a bigger study, and the longitudinal aspect was not analyzed in this study. 17 participants completed all 4 scans; data from 2 participants who were not able to complete all 4 scans due to scheduling constraints was discarded.

#### Freeplay and fNIRS Set-up

The Freeplay set-up was as in Experiment 1. fNIRS set-up was almost identical to that in Experiment 1, with 4 light sources and 8 detectors, to give oxygenation measures in 10 channels on the prefrontal cortex, except with a slightly different sensor layout as depicted in Figure 4.

**Fig. 4.**
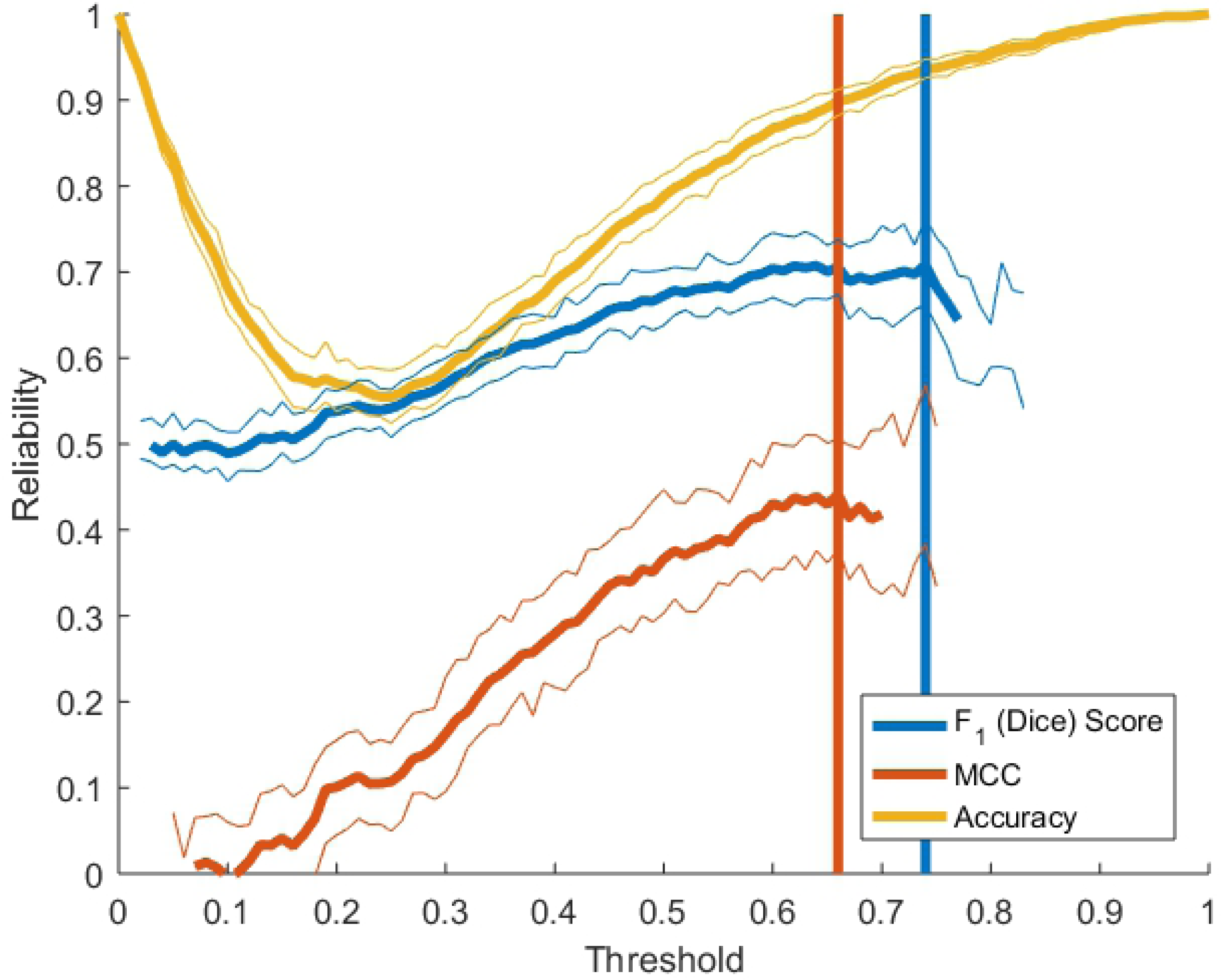
Probe layout for Experiment 2. Sources are in red and detectors are in blue. Channels are in black, labeled 1-5 on the right hemisphere and 6-10 on the left.

#### fNIRS Data Processing

Data was preprocessed and partial correlations computed as in Experiment 1. Again, since there were 10 channels in this probe design, there were 45 (10 choose 2) distinct CPs.

#### Measures of within vs. between subject variance

Kendall’s W and intra-class correlation (ICC) are measures of concordance between groups, frequently used to study test-retest reliability in fMRI data [52–54,17]. In particular, these measures quantify the degree to which scans from the same participant agree compared to scans between different participants. Kendall’s W and ICC both range from 0 to 1, where values approaching 1 indicate high stability of inter-participant variability -- that is, scans are highly reproducible and unique within participants. Smaller values (approaching 0) indicate low stability of inter-participant variability, where scans are highly variable within participants and not differentiable between participants.

Kendall’s *W* and ICC are defined for a given CP as follows. Given ranks (over subjects) of the RSFC in that CP (in each scan), Kendall’s *W* is the mean (over subjects) of the squared deviation of the sum (over scans) of the ranks. That is, if *m* is the number of scans and *n* is the number of subjects, if *r_i,j_* denotes the rank of the *i*^th^ subject’s RSFC in scan *j*, then Kendall’s *W* is defined by the following set of equations:

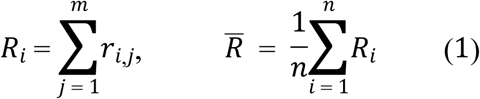

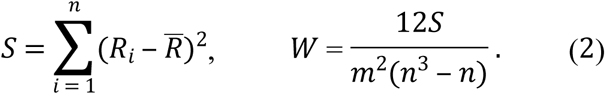

ICC is defined as the proportion of between subjects variation to total variation. That is,

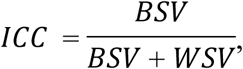

where BSV denotes the between-subjects variance (i.e., the average (over scans) of the variance across subjects) and WSV denotes the within-subjects variance (i.e., the average (over subjects) of the variance across scans).

#### Measures of similarity between binarized connectivity networks

In addition to the partial correlation matrix itself, studies of RSFC are often interested in the network structure of functional connectivity. Studying this requires binarizing the partial correlation matrix (i.e., identifying each CP as either “connected” or “disconnected”. Therefore, to study reliability of the functional connectivity network, we binarized each CP by thresholding the absolute value of its partial correlation value at a “connectivity threshold” θ; absolute values below θ were replaced with 0 (denoting an unconnected CP), and absolute values above θ replaced by 1 (denoting a connected CP).

We then used two indices of inter-scan reliability: the F1 score (a.k.a., Dice coefficient), a general measure of overlap between two sets (twice the ratio of the number of CPs functionally connected in *both* scans to the sum of the numbers of functionally connected CPs over both scans) and the Matthews correlation coefficient (MCC), the correlation between the binarized pattern of 0’s and 1’s, across all 45 CPs. Accuracy (proportion of agreement) between the two binarized scans was not used as a measure of similarity because it is extremely sensitive to the connectivity threshold; for example, using a threshold of 0 (full connectivity) or 1 (no connectivity) results in a perfect accuracy of 1. The raw (continuous, un-thresholded) correlation was also not used, as it is relatively difficult to interpret as a measure of reliability. We chose θ to maximize (over 1000 equally spaced values between 0 and 1) each reliability index (F1 score and MCC) and used LOOCV to obtain an unbiased estimate of each reliability index.

### Results

#### Group-level RSFC correlation

The calculated Pearson and Spearman correlations between mean (across subject) RSFC matrices in each scan (lower triangle of the matrices, excluding main diagonal) were 0.95 with 95% Confidence Interval (CI) (0.88, 0.96) (deoxy-Hb: 0.95 with 95% CI (0.89, 0.95)) and 0.85 with 95% CI (0.64,0.90)(deoxy-Hb: 0.85 with 95% CI (0.58, 0.90)), respectively. This scan1-scan2 correlation for each channel pair is plotted in Figure 5 (panel (c)), along with visualizations of the group level RSFC matrices (panels(a) and (b)). The strong correlation values suggest that conclusions drawn from RSFC group-level matrices should be fairly reproducible. Figure 5(c) <u>also</u> suggests that the presence of adjacent channel pairs, which tend to be strongly functionally connected, skews the RSFC distributions to the right. However, stronger correlations of homologous pairs compared to the rest (non-homologous or adjacent) are still observed. Nevertheless, we investigated the effects of adjacent channel pairs by also running all the analyses with adjacent channels removed – the results are discussed in the section “Post-hoc tests” below.

**Fig. 5.**
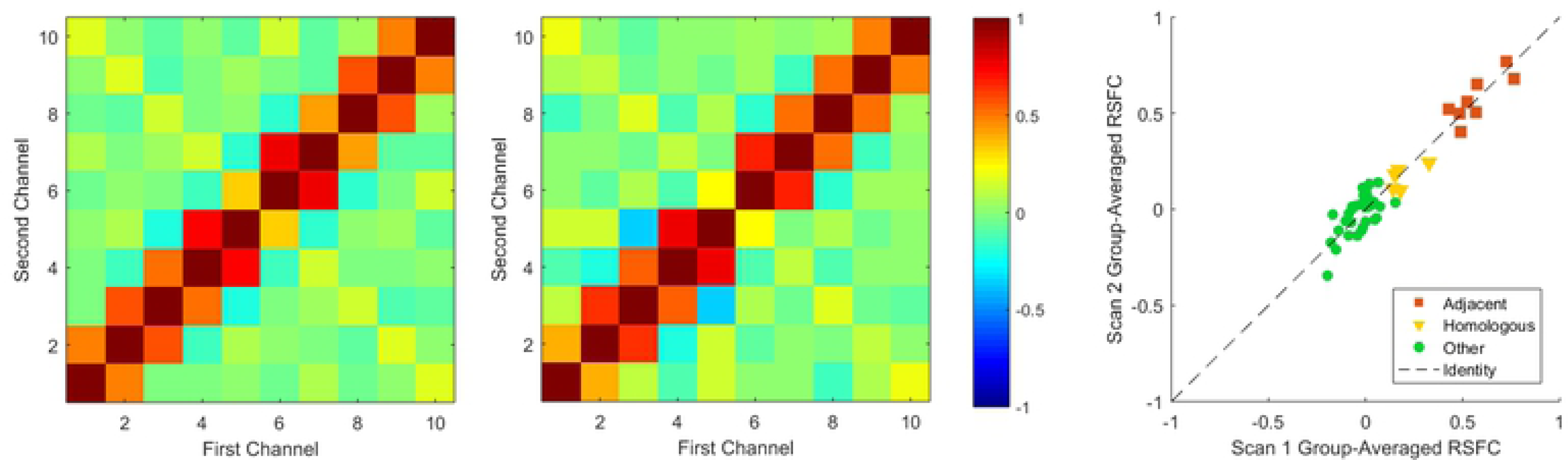
Correlation matrices for scans 1 and 2. Plot of correlation between the two correlation maps shown on the right. Each point corresponds to a CP. Adjacent and homologous CPs, which consistently exhibit stronger functional connectivities, are distinguished from other CPs.

To also measure consistency across scans at the individual level, we calculated Pearson and Spearman correlations between RSFC matrices in each scan, for each subject. The average of those correlations was 0.47 with 95% CI (0.42,0.53) (deoxy-Hb: 0.47 with 95% CI (0.42, 0.54)) for Pearson and 0.43 with 95% CI (0.37, 0.49) (deoxy-Hb: 0.42 with 95% (0.36, 0.48)) for Spearman, suggesting that consistent, detectable signals are present even at the individual level.

#### Within vs. between subject variance

Kendall’s W values (across subjects, between scans, for each CP) had a significantly positive mean (across CPs) of 0.55, with 95% CI (0.52, 0.60) (deoxy-Hb: 0.55 with 95% CI (0.50, 0.59)). Similarly, ICC values (across subjects, between scans), also calculated for each CP, were significantly positive, with a mean of 0.53 with 95% CI (0.51, 0.55) (deoxy-Hb: 0.52 with 95% CI (0.50, 0.55)). Both of these results reflect greater consistency within subjects (or smaller within-subject variance) than between subjects.

#### Consistency of connectivity patterns

The mean (across LOOCV folds) F1 score was 0.71 with 95% CI (0.66, 0.76) (deoxy-Hb: 0.67 with 95% CI (0.63, 0.71)). The mean cutoff threshold chosen was θ=0.73, corresponding to functional connectivity in a mean of 6.4% of CPs. The mean MCC was 0.44 with 95% CI (0.37, 0.51) (deoxy-Hb:0.39 with 95% CI (0.29,0.49)), with a mean cutoff threshold of θ=0.66, corresponding to functional connectivity in a mean of 9.4% of CPs. Within each cross-validation fold, a nested LOOCV was used to select the thresholds that maximized the reliability score. The entire cross-validation curves are shown in Figure 6.

**Fig. 6.**
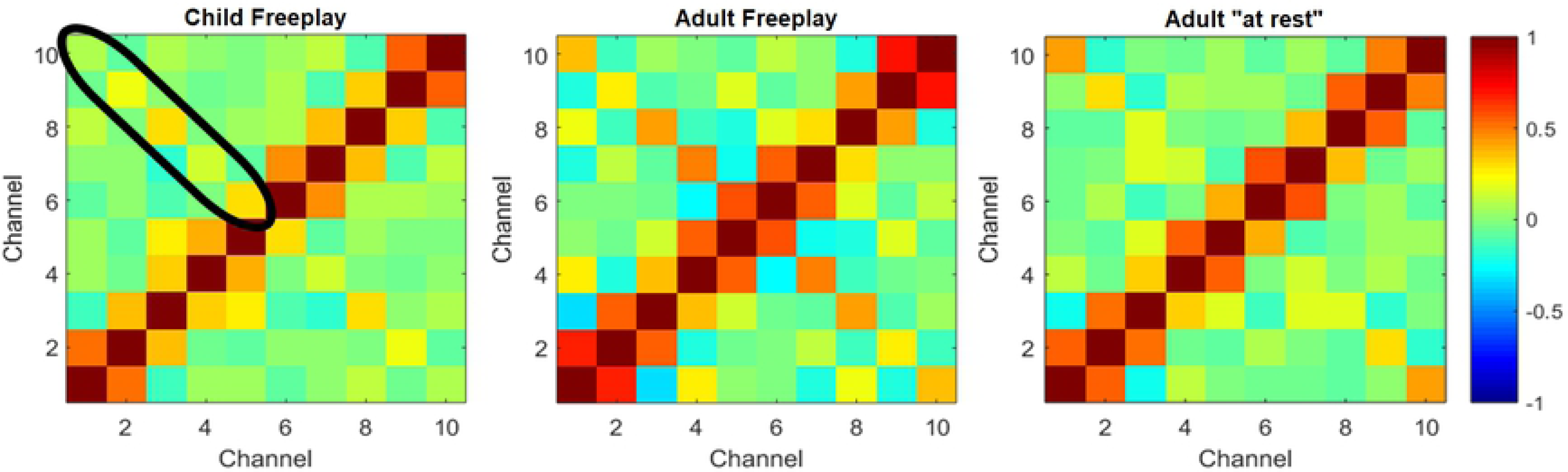
Plot of reliability for each measure over threshold values ranging from 0 to 1. Thin lines show bootstrapped 95% confidence bands. The threshold that maximizes reliability was the same for both F1 and MCC, as indicated by the vertical orange line. For F1 and MCC, reliability is undefined when the entries are all 0 or all 1.

We performed a permutation test comparing the F1 score or MCC to that after randomly permuting the CPs in the second scan for each participant, to test a null model where CPs are randomly identified as functionally connected, with the two scans independent. This test rejected the null for both F1 score and MCC (*p*s<0.01; 1000 permutations).That is, within subjects, we find functional connectivity consistently in the same channels between scans.

Interestingly, similar tests in which second scans were randomly permuted across *participants* (i.e., for a null model of identical/non-distinct participants) were *not* significant (*p*s>0.05), suggesting consistency *across* subjects in Freeplay. This is encouraging from the perspective of trying to identify a common pattern of functional connectivity across individuals. However, further work is needed to understand the sensitivity of the paradigm to individual differences (that may correlate with other quantities of interest, such as age, or behavioral measures), for which we conjecture that the continuous (un-thresholded) CP correlations may be more informative.

#### Post-hoc tests

As is apparent from the group RSFC matrices shown in Figures 2 and 4, adjacent channels are quite strongly correlated with each other -- thus, it is possible that the consistent patterns we detected were driven primarily by these correlations between adjacent channels. To test for this possibility, we re-ran all of the concordance tests after removing adjacent channel pair correlations from the RSFC matrix. The results are shown in Table 1. Although significance levels were generally reduced, all tests that had been significant when adjacent channels were included continued to be significant after their removal, at alpha level 0.05.

**Tab. 1.** Results excluding adjacent channels.

| Quantity | Oxy (95% CI) | Deoxy (95% CI) |
| --- | --- | --- |
| Pearson (Group) | 0.75 (0.32, 0.77) | 0.74 (0.46, 0.78) |
| Spearman (Group) | 0.71 (0.43, 0.75) | 0.71 (0.34, 0.78) |
| Pearson (Subject) | 0.18 (0.09, 0.26) | 0.18 (0.09, 0.25) |
| Spearman (Subject) | 0.17 (0.06, 0.25) | 0.17 (0.07, 0.25) |
| Kendall's W | 0.55 (0.50, 0.61) | 0.55 (0.49, 0.59) |
| ICC | 0.53 (0.49, 0.55) | 0.53 (0.50, 0.56) |
| F1 | 0.51 (0.47, 0.54) | 0.52 (0.49, 0.54) |
| MCC | 0.05 (-0.01, 0.12) | 0.06 (0.00, 0.13) |

## Summary of results

This study explored the feasibility of using fNIRS and Freeplay to measure RSFC in children. Consistent with previous results, we were able to recover connectivity features characteristic of RSFC, helping validate our fNIRS set-up for measuring traditional RSFC. More specifically, homologous channel pairs were significantly more correlated than non-homologous channel pairs, consistent with previous results which found higher coherence between bilateral homologous region pairs than in fronto-posterior pairs or arbitrary pairs in adults [40,41].

Additionally, in adults, correlation patterns in Freeplay were similar to that in traditional resting state -- trained classifiers did not perform significantly better than chance, suggesting that Freeplay may produce a state similar to that in the traditional resting state condition. Since Freeplay was designed to approximate resting state in children, who struggle with the traditional resting state task, this comparison of Freeplay to the traditional task in adult participants serves as an additional check for the viability of fNIRS-Freeplay for measuring RSFC in children. Crucially, all children completed the task and provided usable data, speaking to the practical utility of the Freeplay paradigm for studying RSFC in children. Further, correlations in adults and children in Freeplay showed different patterns, from which a trained classifier was able to predict “adult” or “child” with high accuracy -- this is in line with our expectations given that we know RSNs develop significantly with age. Finally, Experiment 2 demonstrated inter-scan reliability, in that similar connectivity patterns are found between 2 independent scans of the same individual, and that within-subject variance is significantly lower than between-subject variance. This reliability is observed for both the raw RSFC matrices and the resulting connectivity networks after thresholding appropriately, and is observed both with and without adjacent channel pairs.

## Limitations and future directions

The test-retest reliability value for ICC (0.53) was lower in our paper than in a previous paper (0.7) by Niu et al. looking at RSFC test-retest reliability in adults [19]. Some possible reasons for this may be that: 1) since we are measuring in children, the data may be noisier, and 2) they measured functional connectivity in terms of Pearson (rather than partial) correlations, which can be inflated by other sources of correlation between channels besides functional brain correlations.

Although our study compared “Freeplay” to the traditional “at rest” condition measured with adults, it did not compare to either a true task condition, or a “movie-watching” condition. Comparing the functional connectivity patterns in “Freeplay” with not just “at rest” but also these other conditions may help us better characterize RSFC. For example, if we are able to show that a trained classifier can effectively distinguish between adults in the “at rest” or “Freeplay” conditions from those in the movie watching condition, it would provide a comparison condition for “Freeplay” as well as corroborate previous findings that “movie watching” alters resting state.

The current study introduced a new paradigm for measuring resting state functional connectivity in children that combines fNIRS and a method called Freeplay to help increase participant compliance and induce a state similar to adult resting state in children. We demonstrated the practical feasibility as well as the construct validity of the Freeplay-fNIRS paradigm for studying RSFC in children by measuring correlations between pairs of regions in the PFC. As previously discussed, the PFC is a central player in the central executive network (CEN), and studying its RSFC patterns over development is crucial to understanding how the CEN develops and supports executive function (EF). Zhao et al. [55] recently found that network properties of RSFC in the PFC, as measured by fNIRS, were correlated with varying performance in EF tasks in adults. An important next step will be to extend those investigations to children. Future work can explore the changes in PFC connectivity structures that parallel observed cognitive development in EF or other domains subserved by the PFC. As a practical setup for measuring RSFC in children, the fNIRS-Freeplay paradigm will allow investigation of questions such as these to advance understanding of the neural mechanisms of cognitive development.

## Acknowledgements

We thank the children, parents, and teachers of the CMU Children’s School for making this work possible.

## References

[1] Moussa MN, Steen MR, Laurienti PJ, Hayasaka S. Consistency of network modules in resting-state FMRI connectome data. PloS one. 2012 Aug 31;7(8):e44428.

[2] Friston KJ. Functional and effective connectivity in neuroimaging: a synthesis. Human brain mapping. 1994;2(1-2):56–78.

[3] Rosazza, C., & Minati, L. (2011). Resting-state brain networks: literature review and clinical applications. Neurological Sciences, 32(5), 773–785.

[4] Fox MD, Raichle ME. Spontaneous fluctuations in brain activity observed with functional magnetic resonance imaging. Nature reviews neuroscience. 2007 Sep;8(9):700.

[5] Biswal, B., Zerrin Yetkin, F., Haughton, V. M., & Hyde, J. S. (1995). Functional connectivity in the motor cortex of resting human brain using echo‐planar mri. Magnetic resonance in medicine, 34(4), 537–541.

[6] Cordes, D., Haughton, V. M., Arfanakis, K., Wendt, G. J., Turski, P. A., Moritz, C. H., … & Meyerand, M. E. (2000). Mapping functionally related regions of brain with functional connectivity MR imaging. American Journal of Neuroradiology, 21(9), 1636–1644.

[7] Seeley, W. W., Menon, V., Schatzberg, A. F., Keller, J., Glover, G. H., Kenna, H., … & Greicius, M. D. (2007). Dissociable intrinsic connectivity networks for salience processing and executive control. Journal of Neuroscience, 27(9), 2349–2356.

[8] Stevens, W. D., Buckner, R. L., & Schacter, D. L. (2009). Correlated low-frequency BOLD fluctuations in the resting human brain are modulated by recent experience in category-preferential visual regions. Cerebral cortex, 20(8), 1997–2006.

[9] Jolles DD, van Buchem MA, Crone EA, Rombouts SA. A comprehensive study of whole-brain functional connectivity in children and young adults. Cerebral cortex. 2010 Jun 11;21(2):385–91.

[10] Barkhof F, Haller S, Rombouts SA. Resting-state functional MR imaging: a new window to the brain. Radiology. 2014 Jul;272(1):29–49.

[11] Supekar K, Musen M, Menon V. Development of large-scale functional brain networks in children. PLoS biology. 2009 Jul 21;7(7):e1000157.

[12] Fair DA, Cohen AL, Power JD, Dosenbach NU, Church JA, Miezin FM, Schlaggar BL, Petersen SE. Functional brain networks develop from a “local to distributed” organization. PLoS computational biology. 2009 May 1;5(5):e1000381.

[13] Byars AW, Holland SK, Strawsburg RH, Bommer W, Dunn RS, et al. (2002): Practical aspects of conducting large-scale functional magnetic resonance imaging studies in children. J Child Neurol 17:885–889.

[14] Yerys BE, Jankowski KF, Shook D, Rosenberger LR, Barnes KA, Berl MM, Ritzl EK, VanMeter J, Vaidya CJ, Gaillard WD. The fMRI success rate of children and adolescents: typical development, epilepsy, attention deficit/hyperactivity disorder, and autism spectrum disorders. Human brain mapping. 2009 Oct;30(10):3426–35.

[15] Betti V, Della Penna S, de Pasquale F, Mantini D, Marzetti L, Romani GL, Corbetta M. Natural scenes viewing alters the dynamics of functional connectivity in the human brain. Neuron. 2013 Aug 21;79(4):782–97.

[16] Uddin LQ, Supekar K, Menon V. Typical and atypical development of functional human brain networks: insights from resting-state FMRI. Frontiers in systems neuroscience. 2010 May 21;4:21.

[17] Thomason ME, Dennis EL, Joshi AA, Joshi SH, Dinov ID, Chang C, Henry ML, Johnson RF, Thompson PM, Toga AW, Glover GH. Resting-state fMRI can reliably map neural networks in children. Neuroimage. 2011 Mar 1;55(1):165–75.

[18] Braun U, Plichta MM, Esslinger C, Sauer C, Haddad L, Grimm O, Mier D, Mohnke S, Heinz A, Erk S, Walter H. Test–retest reliability of resting-state connectivity network characteristics using fMRI and graph theoretical measures. Neuroimage. 2012 Jan 16;59(2):1404–12.

[19] Niu H, Li Z, Liao X, Wang J, Zhao T, Shu N, Zhao X, He Y. Test-retest reliability of graph metrics in functional brain networks: a resting-state fNIRS study. PLoS One. 2013 Sep 9;8(9):e72425.

[20] Geng S, Liu X, Biswal BB, Niu H. Effect of resting-state fNIRS scanning duration on functional brain connectivity and graph theory metrics of brain network. Frontiers in neuroscience. 2017 Jul 20;11:392.

[21] Raschle N, Zuk J, Ortiz-Mantilla S, Sliva DD, Franceschi A, Grant PE, Benasich AA, Gaab N. Pediatric neuroimaging in early childhood and infancy: challenges and practical guidelines. Annals of the New York Academy of Sciences. 2012 Apr;1252:43.

[22] Power JD, Barnes KA, Snyder AZ, Schlaggar BL, Petersen SE. Spurious but systematic correlations in functional connectivity MRI networks arise from subject motion. Neuroimage. 2012 Feb 1;59(3):2142–54.

[23] Nishiyori R. fNIRS: An emergent method to document functional cortical activity during infant movements. Frontiers in psychology. 2016 Apr 20;7:533.

[24] Patriat R, Molloy EK, Meier TB, Kirk GR, Nair VA, Meyerand ME, Prabhakaran V, Birn RM. The effect of resting condition on resting-state fMRI reliability and consistency: a comparison between resting with eyes open, closed, and fixated. Neuroimage. 2013 Sep 1;78:463–73.

[25] Liu WC, Flax JF, Guise KG, Sukul V, Benasich AA. Functional connectivity of the sensorimotor area in naturally sleeping infants. Brain research. 2008 Aug 5;1223:42–9.

[26] Horovitz SG, Fukunaga M, de Zwart JA, van Gelderen P, Fulton SC, Balkin TJ, Duyn JH. Low frequency BOLD fluctuations during resting wakefulness and light sleep: A simultaneous EEG‐fMRI study. Human brain mapping. 2008 Jun;29(6):671–82.

[27] Horovitz SG, Braun AR, Carr WS, Picchioni D, Balkin TJ, Fukunaga M, Duyn JH. Decoupling of the brain’s default mode network during deep sleep. Proceedings of the National Academy of Sciences. 2009 Jul 7;106(27):11376–81.

[28] Boly M, Perlbarg V, Marrelec G, Schabus M, Laureys S, Doyon J, Pélégrini-Issac M, Maquet P, Benali H. Hierarchical clustering of brain activity during human nonrapid eye movement sleep. Proceedings of the National Academy of Sciences. 2012 Apr 10;109(15):5856–61.

[29] Spoormaker VI, Gleiser P, Czisch M. Frontoparietal connectivity and hierarchical structure of the brain’s functional network during sleep. Frontiers in neurology. 2012 May 17;3:80.

[30] de Bie HM, Boersma M, Wattjes MP, Adriaanse S, Vermeulen RJ, Oostrom KJ, Huisman J, Veltman DJ, Delemarre-Van de Waal HA. Preparing children with a mock scanner training protocol results in high quality structural and functional MRI scans. European journal of pediatrics. 2010 Sep 1;169(9):1079–85.

[31] Greene DJ, Koller JM, Hampton JM, Wesevich V, Van AN, Nguyen AL, Hoyt CR, McIntyre L, Earl EA, Klein RL, Shimony JS. Behavioral interventions for reducing head motion during MRI scans in children. NeuroImage. 2018 May 1;171:234–45.

[32] Barker JW, Aarabi A, Huppert TJ. Autoregressive model based algorithm for correcting motion and serially correlated errors in fNIRS. Biomedical optics express. 2013 Aug 1;4(8):1366–79.

[33] Emerson RW, Short SJ, Lin W, Gilmore JH, Gao W. Network-level connectivity dynamics of movie watching in 6-year-old children. Frontiers in human neuroscience. 2015 Nov 23;9:631.

[34] Cantlon JF, Li R. Neural activity during natural viewing of Sesame Street statistically predicts test scores in early childhood. PLoS biology. 2013 Jan 3;11(1):e1001462.

[35] Vanderwal T, Kelly C, Eilbott J, Mayes LC, Castellanos FX. Inscapes: A movie paradigm to improve compliance in functional magnetic resonance imaging. Neuroimage. 2015 Nov 15;122:222–32.

[36] Casey BJ, Giedd JN, Thomas KM. Structural and functional brain development and its relation to cognitive development. Biological psychology. 2000 Oct 1;54(1-3):241–57.

[37] Diamond A. The early development of executive functions. Lifespan cognition: Mechanisms of change. 2006;210:70–95.

[38] Diamond A. Executive functions. Annual review of psychology. 2013 Jan 3;64:135–68.

[39] Lin HY, Tseng WY, Lai MC, Matsuo K, Gau SS. Altered resting-state frontoparietal control network in children with attention-deficit/hyperactivity disorder. Journal of the International Neuropsychological Society. 2015 Apr;21(4):271–84.

[40] Sasai S, Homae F, Watanabe H, Taga G. Frequency-specific functional connectivity in the brain during resting state revealed by NIRS. Neuroimage. 2011 May 1;56(1):252–7.

[41] Sasai S, Homae F, Watanabe H, Sasaki AT, Tanabe HC, Sadato N, Taga G. A NIRS–fMRI study of resting state network. Neuroimage. 2012 Oct 15;63(1):179–93.

[42] Wang J, Dong Q, Niu H. The minimum resting-state fNIRS imaging duration for accurate and stable mapping of brain connectivity network in children. Scientific reports. 2017 Jul 25;7(1):6461.

[43] Abdelnour AF, Huppert T. Real-time imaging of human brain function by near-infrared spectroscopy using an adaptive general linear model. Neuroimage. 2009 May 15;46(1):133–43.

[44] Huppert TJ. History of diffuse optical spectroscopy of human tissue. InOptical Methods and Instrumentation in Brain Imaging and Therapy 2013 (pp. 23–56). Springer, New York, NY.

[45] Whiteman AC, Santosa H, Chen DF, Perlman SB, Huppert T. Investigation of the sensitivity of functional near-infrared spectroscopy brain imaging to anatomical variations in 5-to 11-year-old children. Neurophotonics. 2017 Sep;5(1):011009.

[46] Cui X, Bray S, Reiss AL. Functional near infrared spectroscopy (NIRS) signal improvement based on negative correlation between oxygenated and deoxygenated hemoglobin dynamics. Neuroimage. 2010 Feb 15;49(4):3039–46.

[47] Chiarelli AM, Maclin EL, Fabiani M, Gratton G. A kurtosis-based wavelet algorithm for motion artifact correction of fNIRS data. Neuroimage. 2015 May 15;112:128–37.

[48] Sakakibara E, Homae F, Kawasaki S, Nishimura Y, Takizawa R, Koike S, Kinoshita A, Sakurada H, Yamagishi M, Nishimura F, Yoshikawa A. Detection of resting state functional connectivity using partial correlation analysis: A study using multi-distance and whole-head probe near-infrared spectroscopy. NeuroImage. 2016 Nov 15;142:590–601.

[49] Marrelec G, Krainik A, Duffau H, Pélégrini-Issac M, Lehéricy S, Doyon J, Benali H. Partial correlation for functional brain interactivity investigation in functional MRI. Neuroimage. 2006 Aug 1;32(1):228–37.

[50] Tibshirani R. Regression shrinkage and selection via the lasso. Journal of the Royal Statistical Society: Series B (Methodological). 1996 Jan;58(1):267–88.

[51] Cunningham P, Delany SJ. k-Nearest neighbour classifiers. Multiple Classifier Systems. 2007 Mar 27;34(8):1–7.

[52] Caceres A, Hall DL, Zelaya FO, Williams SC, Mehta MA. Measuring fMRI reliability with the intra-class correlation coefficient. Neuroimage. 2009 Apr 15;45(3):758–68.

[53] Meltzer JA, Postman-Caucheteux WA, McArdle JJ, Braun AR. Strategies for longitudinal neuroimaging studies of overt language production. Neuroimage. 2009 Aug 15;47(2):745–55.

[54] Shehzad Z, Kelly AC, Reiss PT, Gee DG, Gotimer K, Uddin LQ, Lee SH, Margulies DS, Roy AK, Biswal BB, Petkova E. The resting brain: unconstrained yet reliable. Cerebral cortex. 2009 Feb 16;19(10):2209–29.

[55] Zhao J, Liu J, Jiang X, Zhou G, Chen G, Ding XP, Fu G, Lee K. Linking resting-state networks in the prefrontal cortex to executive function: a functional near infrared spectroscopy study. Frontiers in neuroscience. 2016 Oct 7;10:452.

